# Five years of land surface phenology in a large-scale flooding and draining manipulation in a coastal Arctic ecosystem

**DOI:** 10.1101/146662

**Authors:** Santonu Goswami, John A. Gamon, Sergio Vargas, Craig E. Tweedie

**Affiliations:** Climate Change Science Institute and Environmental Sciences Division, Oak Ridge National Laboratory, Oak Ridge, TN, USA 3783 1.; Dept. of Earth and Atmospheric Sciences, 1-26 Earth Sciences Building, University of Alberta, Edmonton, Alberta, Canada T6G 2E3.; Environmental Science and Engineering Program, University of Texas at El Paso, El Paso, Texas, USA 79968.

## Abstract

This study was motivated by the knowledge gap for observing the complex interplay between surface hydrology and plant phenology in arctic landscapes and was conducted as part of a large scale, multi investigator flooding and draining experiment near Barrow, Alaska (71°17’01” N, 156°35’48” W) during 2005 - 2009. Hyperspectral reflectance data were collected in the visible to near IR region of the spectrum using a robotic tram system that operated along a 300m transects during the snow free growing period between June and August, 2005-09. Interannual patterns of land-surface phenology (NDVI) unexpectedly lacked marked differences under experimental conditions. Measurement of NDVI was, however, compromised for presence of surface water. Land-surface phenology and surface water was negatively correlated, which held when scaled to a 2km by 2km MODIS subset of the study area. This result suggested that published findings of ‘greening of the Arctic’ may relate to a ‘drying of the Arctic’ i.e. reduced surface water in vegetated high-latitude landscapes where surface water is close to ground level.

## Introduction

Detecting the biotic responses of arctic terrestrial ecosystems to environmental change is essential for understanding the consequences of global change and the future state of the Arctic and Earth Systems (McGuire *et al.*, 2009, Sitch *et al.*, 2007). Many observed and modeled change climate change responses of arctic terrestrial ecosystems such as surface energy budgets (Chapin *et al.*, 2005, Euskirchen *et al.*, 2007) land-atmosphere carbon exchange (Merbold *et al.*, 2009, Wolf *et al.*, 2008) geomorphic processes (Lawrence & Slater, 2005, McNamara & Kane, 2009) provision of ecosystem goods and services plant and landscape phenology (Bhatt *et al.*, 2010, Jia *et al.*, 2009, Walker *et al.*, 2006) and response to warming (Arft *et al.*, 1999, Walker *et al.*, 2006) are related to surface hydrology. Of particular concern, is how the combined alteration of air and soil temperature, and altered surface hydrology will affect the structure and function of arctic terrestrial ecosystems, particularly ecosystem carbon balance (Post *et al.*, 2009, Schuur *et al.*, 2009). If the biomass increases forecast for most arctic terrestrial ecosystems (Euskirchen *et al.*, 2009, Kimball *et al.*, 2007) do not offset predicted losses of greenhouse gases to the atmosphere due to permafrost thaw and microbial decomposition of the substantial arctic soil carbon store (Tarnocai *et al.*, 2009), greenhouse warming will be enhanced (Kimball *et al.*, 2006).

Several key studies have shown that plants can serve as effective indicators of environmental change (Arft *et al.*, 1999, Walker *et al.*, 2006). At the leaf and plant level, plants can respond to change with an increase or decrease in green-leaf biomass (Riedel *et al.*, 2005), pigments, and leaf water content (Hudson, 2010). At the plant community and ecosystem level, plant responses to change include shifts in species composition and abundance and ecosystem processes such as nutrient cycling (Johnson *et al.*, 2002) and surface energy balance (Chapin *et al.*, 2005, Euskirchen *et al.*, 2007). All of these multilevel plant responses to change can be observed and quantified using optical remote sensing techniques (La Puma *et al.*, 2007, Olthof *et al.*, 2008). In the Arctic however, there has been a general paucity of ground based studies examining plant responses to change using remote sensing approaches (Boelman *et al.*, 2003). As such, the transferability of algorithms developed for detecting these phenomenon in other ecosystems remain poorly tested and few studies have documented seasonal and inter-annual dynamics of land surface properties using these ground based methods. These methods have, however provided cost effective ground to satellite scaling (Ueyama *et al.*, 2009) of leaf to landscape ecosystem properties and processes in other ecosystems (Hilker *et al.*, 2008) and could help to overcome many of the deficiencies associated with satellite remote sensing in the Arctic (Stow *et al.*, 2004).

Land surface phenology (LSP) is a measure of the seasonal variation in vegetated land surfaces that occurs in response to climate and can be observed using ground to satellite remote sensing approaches **(Friedl et al. 2006).** Using suitable vegetation spectral indices such as the NDVI (Normalized Difference Vegetation Index), which has been shown to be useful to estimate above ground green biomass and plant cover (Boelman *et al.*, 2003, Epstein *et al.*, 2008, Gamon *et al.*, 1995) changes in the timing and extent of peak greenness (de Beurs & Henebry, 2010), rates of green-up and senescence (Stow *et al.*, 1993), differences between different landscape units, and change over time (Bhatt *et al.* 2010) can be quantified. NDVI is affected by various environmental factors (Galvao *et al.*, 2004, Matsushita *et al.*, 2007, Nagol *et al.*, 2009). Nonetheless, if executed correctly, landscape phenology can serve as an indicator of ecosystem state and change over time.

This study aims to improve our understanding of the spatial and temporal dynamics and variability in tundra land-surface phenology in a hydrological manipulation experiment. The study further aims to understand how land-surface phenology is affected by altered surface hydrology, and if such alterations can be documented with optical remote sensing at both the ground and satellite scales. Specifically, we examine inter-annual variability in landscape phenology and how changes in surface hydrology conducted in association with a large scale flooding and draining experiment affected seasonal dynamics of land-surface phenology. Building on a previous study that showed that Normalized Difference Surface Water Index (NDSWI) could be used to characterize surface water in the study area (Goswami *et al.*, 2011), we also explore how NDVI dynamics relate to those of NDSWI and if these dynamics scale to the satellite scale using MODIS time series surface reflectance data obtained for the study site over the snow free growing period between 2000 and 2010.

## Methods

### Study Site and Experimental Design

This study was conducted within the Biocomplexity flooding and draining experiment located on Barrow Environmental Observatory (BEO) near Barrow, Alaska, 71°17’01” N, 156°35’48” W. This large-scale experimental infrastructure was designed to examine the role of altered water table on land - atmosphere carbon, water and energy balance. After three years of baseline data collection from 2005 – 2007 (2007 included a failed 2-week long experimental manipulation at the beginning of the snow free period), three experimental treatments were initiated and sustained through the 2008 and 2009 snow free periods. Treatments consisted of a control section that was unmanipulated and held relative to water tables outside the experimental area, and flooded and drained sections where the water table was raised and lowered by 10 cm relative to the control treatment respectively (Fig. 1.2b). This experimental design explicitly allowed for inter-annual variability to be incorporated, which in this unreplicated experiment was essential for assessing the integrity of the experiment by testing the hypothesis that during a wet year, ecosystem properties and processes in the control treatment would be similar to those of the wet treatment area in a dry year.

**Figure 1.**
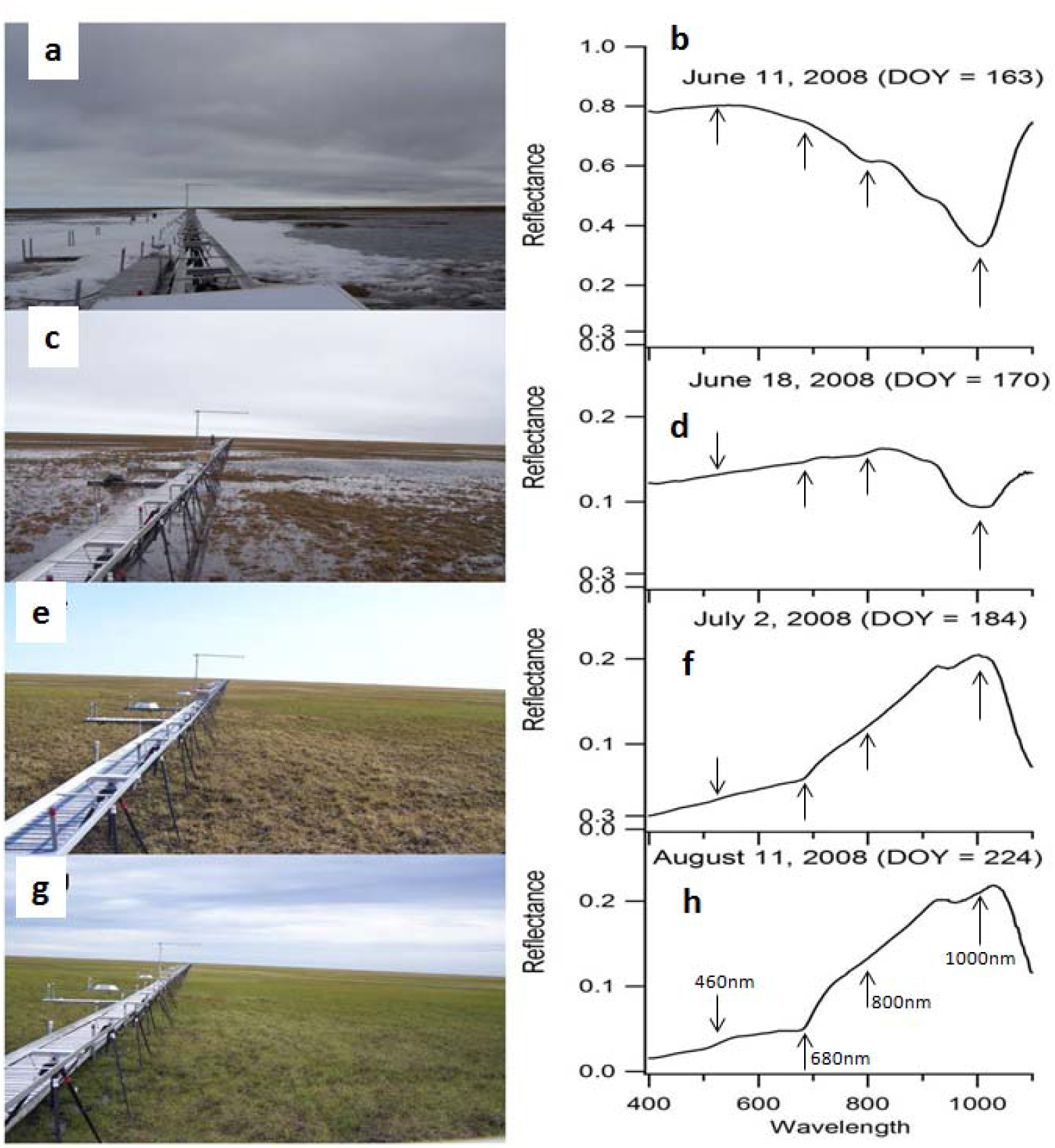
Seasonal changes in land-surface phenology visualized on the left with photographs and on the right with reflectance collected on the same day as the photographs and averaged over the 300m length of the south tramline. Surface spectral reflectance was able to capture the transition from snow melt to maximum greenness in the study area as shown above. The first photograph/plot pair shows the spectral characteristics of melting snow (1a, 1b), followed by the spectral characteristics of the experimental area immediately after snowmelt (1c, 1d), in the initial growth period (1e, 1f), and during the peak growing period (1g, 1h). The arrows pointing at 680nm and 800nm show the wavelengths used for calculating NDVI and the at 460nm and 100nm show the wavelengths used for calculating NDSWI.

**Figure 2.**
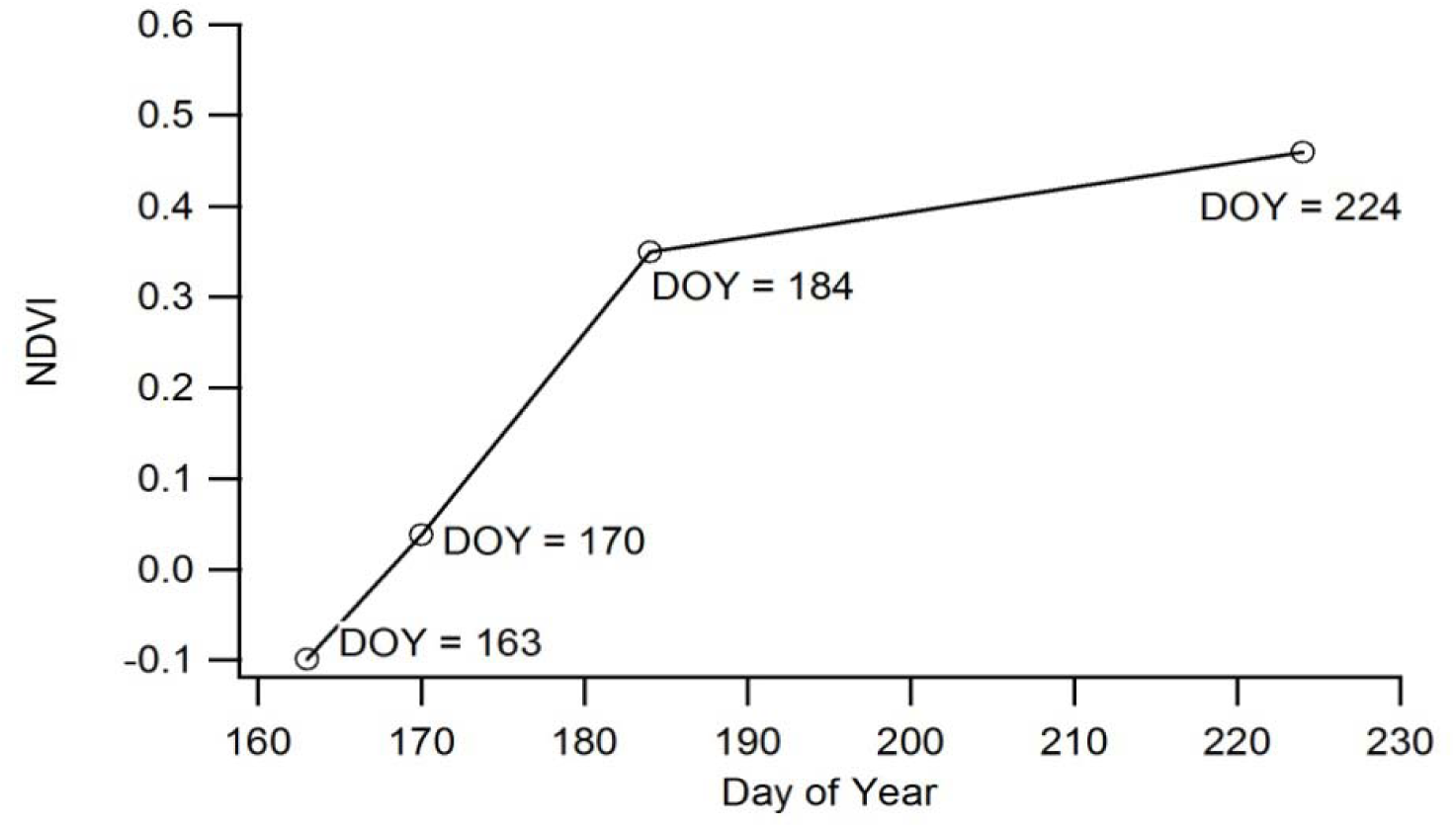
Change in land-surface phenology (NDVI) at different days of the year (DOY) over the course of the 2008 snow free period for the same dates shown in figure 3.1.

Vegetation within the Biocomplexity experimental area is predominantly moist and wet graminoid tundra (Teh *et al.*, 2009). The study site is underlain by continuous ice and organic carbon-rich permafrost and the general landscape has characteristic thermokarst features common on the Arctic Coastal Plain including shallow ponds, high and low-centered polygons, polygon rims and troughs, and meadows **(Brown et al. 1980**). Active layer depth is typically less than 50 centimeters in the experimental area (Shiklomanov *et al.*, 2010). Soils are generally moisture rich due to shallow drainage gradients and impermeable continuous permafrost, and are predominantly comprised of cryoturbated gelisols (Bockheim *et al.*, 1999). Rates of evapotranspiration are relatively low (Liljedahl *et al.*, 2011).

The study area has characteristic long and cold winters with a mean winter air temperature of -26.6°C in February. Summers are relatively cool and wet and mean maximum temperatures in July typically reach 4.7°C (Bockheim *et al.*, 1999). During summer, 24 hour daylight prevails for 85 days between May 10 and August 2. The mean annual temperature for Barrow is approximately -12.0°C (Oberbauer *et al.*, 2007). Snowmelt occurs in early June when average daily air temperatures rise above 0°C, and snow typically begins to accumulate in mid to late September (Hobbie, 1975).

### Reflectance Measurements

A robotic tram system (Gamon *et al.*, 2006) was used to collect the field reflectance data used for this study. The tram system consisted of three 300 m long transects (“tramlines”) with one tramline located in each of the three treatment areas and oriented in an east-west direction spanning the entire width of the lakebed. Each of the tramlines was 200 meters in length in 2005 and was extended by 100 meters in early spring of 2006. This required snow removal for the length of the extension (meter 201 – meter 300) for each of the tramlines. This infrastructure provided an ideal research platform for repeat measurement of surface spectral properties of the same study area affected by contrasting surface hydrology regimes throughout the snow-free period. A detailed description of the experimental infrastructure used for this study is given in (Goswami *et al.*, 2011).

Reflectance data were collected at every meter along each tramline using a dual detector Unispec DC field spectrometer (PP Systems, Amesbury, MA, USA) housed within a robotic cart that was operated semi-autonomously (Gamon *et al.*, 2006). The cart travelled along the tramline at an average speed of 20cm/s, taking approximately 25 minutes to travel the full length of the 300m long tramline. The spectrometer was activated when a mechanical switch mounted on the base of the robotic cart was triggered as the cart passed over crossbars positioned every meter along each tramline. One run along the length of one tramline produced three hundred spectral measurements every time the robotic cart was operated. A footprint area of approximately 1m^2^ was achieved by positioning the downward looking foreoptic (with a 20 degrees field of view) on a south-facing boom mounted to the cart approximately 3 m above ground level. The upward and downward looking detectors of the field spectrometer were cross calibrated in the field using a white reflectance panel with 99% reflectance (Spectralon, labsphere, North Sutton, NH, USA) at the beginning and at the end of each run as described in Gamon *et al*. (2006). The Unispec DC had a nominal range of operation between 303 and 1148 nm in 256 contiguous bands with a spectral resolution of approximately 3 nm and a full width at half maximum of approximately 10 nm. Tram measurements were made three times every week during the field season if weather permitted (no extreme winds, fog, rain, or falling snow). Data used for the study documented in this paper was extracted from datasets that spanned the majority of the snow free growing period between mid June and mid August in all years between 2005 and 2009.

This study utilizes the Normalized difference vegetation index (NDVI) to document seasonal and inter-annual landscape phenological dynamics. NDVI values were calculated for each sampled meter of each tramline for every day measurements were made (n=300 measurements/tram/day) using the following equation (Gamon et al. 2006):

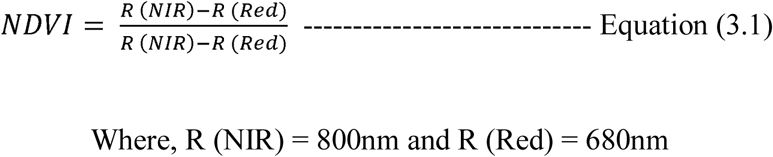

To visualize seasonal trends in NDVI, mean NDVI for each tramline and sample time was calculated for each of the five years of measurement (2005 – 2009).

Additionally, Normalized Difference Surface Water Index (NDSWI) (Goswami *et al.* 2011) was calculated to investigate the effect of surface water on reflectance spectra as follows:

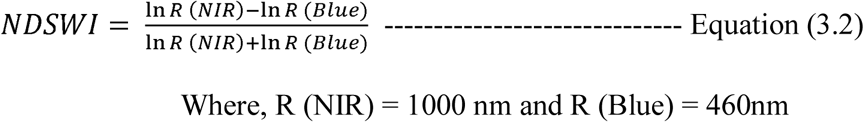

### Ancillary Measurements

A range of ancillary measurements were made in addition to the reflectance measurements described above. Water table depth data were collected manually every 10 meters along each tramline within a day of the spectral measurements being acquired. Water table depth measurements were made relative to the ground surface using perforated PVC tubes inserted in holes drilled to below the active layer. Negative numbers indicate water levels below the ground surface whereas positive numbers indicate pooling of water above the ground surface.

Digital photos of each of the tramline footprints were taken several times during the 2006-2009 field seasons using a Nikon Coolpix 5400 camera mounted on the boom of the robotic tram looking downward. The camera was triggered manually every meter of the tramline using a remote trigger so that the footprint of the spectral measurements was captured. Oblique landscape photographs of each tramline area were also taken from the west end of each tramline once every week from early June to the end of August in 2009.

A HOBO automated weather station (AWS) was used to measure air temperature, relative humidity, precipitation and photosynthetic active radiation (PAR) between early June and late August at one minute intervals. The AWS was installed at the west end of the central tramline. Surface elevations at every meter of each tramline were measured using a survey grade Trimble differential GPS unit to document microtopographic variability along each of the tramlines.

### Statistical Analysis

To determine the effect of experimental flooding and draining on seasonal LSP, a series of regression analyses were used. For pre-treatment years, regression models were built to predict mean NDVI values of the north and central tramlines from the measured NDVI at the southern tramline. During experimental years (2008-2009), to quantify treatment effects, measured NDVI from the north and central tramlines was subtracted from NDVI predicted with the use of the regression equations that modeled pre-treatment conditions The extent to which treatment effects could be determined for 2009 is limited by an interruption of water levels in the control treatment late in the growing season, which were experimentally altered by other investigators as part of a short-term experiment flooding experiment during the middle two weeks of August.

A gamut of interacting factors is known to affect both the vegetation signal and spectral measurement of NDVI in tundra landscapes (Hope *et al.*, 1999, Jia *et al.*, 2003, Stow *et al.*, 2004). To quantitatively explore the environmental controls of seasonal NDVI at the study site, regression tree analysis was performed using JPM 7.0 (SAS Institute, Inc. Cary NC) statistical software where NDVI data for 2005 – 2009 averaged for each tramline was included as the y-response (dependent variable). X-response variables (independent variables) included day since snow melt (DSSM), mean daily air temperature, water table depth, and total daily PAR. Daily average temperature, water table depth and daily total PAR were calculated from AWS measurements. Regression tree analysis is a practical and informative (Kheir *et al.*, 2008) procedure useful for data exploration and several studies have used this approach to explore the presence of ecological thresholds (Kheir *et al.*, 2008, Sickman *et al.*, 2002) **Michaelson et al. 2004, Franklin 1998, and Kandrika 2008**).

NDSWI was developed to characterize the cover and depth of surface water from spectral bands common to a range of ground, aerial, and satellite optical remote sensing instruments, and increases with increasing water cover and depth (Goswami *et al.* 2011). For this study, NDWSI was calculated following Equation (3.2) for each year/tramline/measurement combination. Time series of mean NDSWI and NDVI values were plotted over the snow free period of the growing season for all five years of study (2005 – 2009) to explore how a change in NDVI correlated with a change in NDSWI (i.e. surface water) **(Fig. 3.9).** Differences in NDVI and NDSWI values between sampling times within a season were first calculated for a given measurement day by subtracting the NDVI and NDSWI value from the previous sampling day (e.g. “day 175” – “day 173”, “day 177” – “day 175”, “day 179” – “day 177” etc.) for all three treatment areas and treatment years (i.e. 2008 and 2009). Following this, linear regression models were developed between “the change in NDVI within a given sampling period” and “change in NDSWI within a given sampling period” for both the flooded (north), drained (central), and control (south) treatment areas. The drained and control sections of the study area had very little surface water present compared to the flooded section during treatment years so regression models were developed separately for the flooded section.

**Figure 3.**
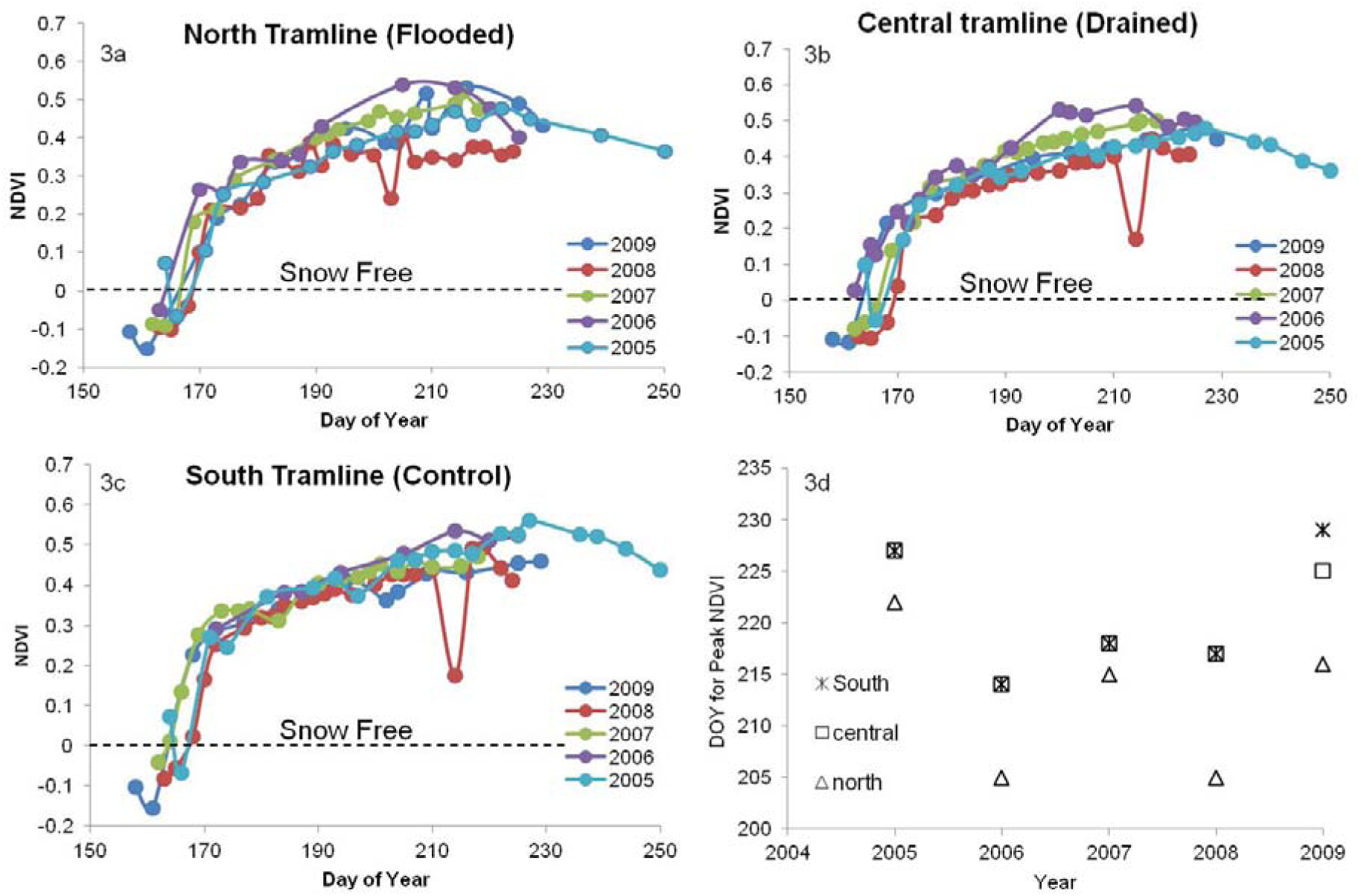
Seasonal and inter-annual patterns of land-surface phenology (mean NDVI for all measurements along each tramline) for each of the treatment areas for pre-treatment (2005, 2006, 2007) and treatment (2008, 2009) years. Initiation of greening was dependent on snow melt and a moderate level of variability in NDVI was noted for most treatments and years. The DOY corresponding to peak season NDVI is shown in 3d. The standard deviations for snow-free periods for all the tramlines for all the years were 0.1.

**Figure 4.**
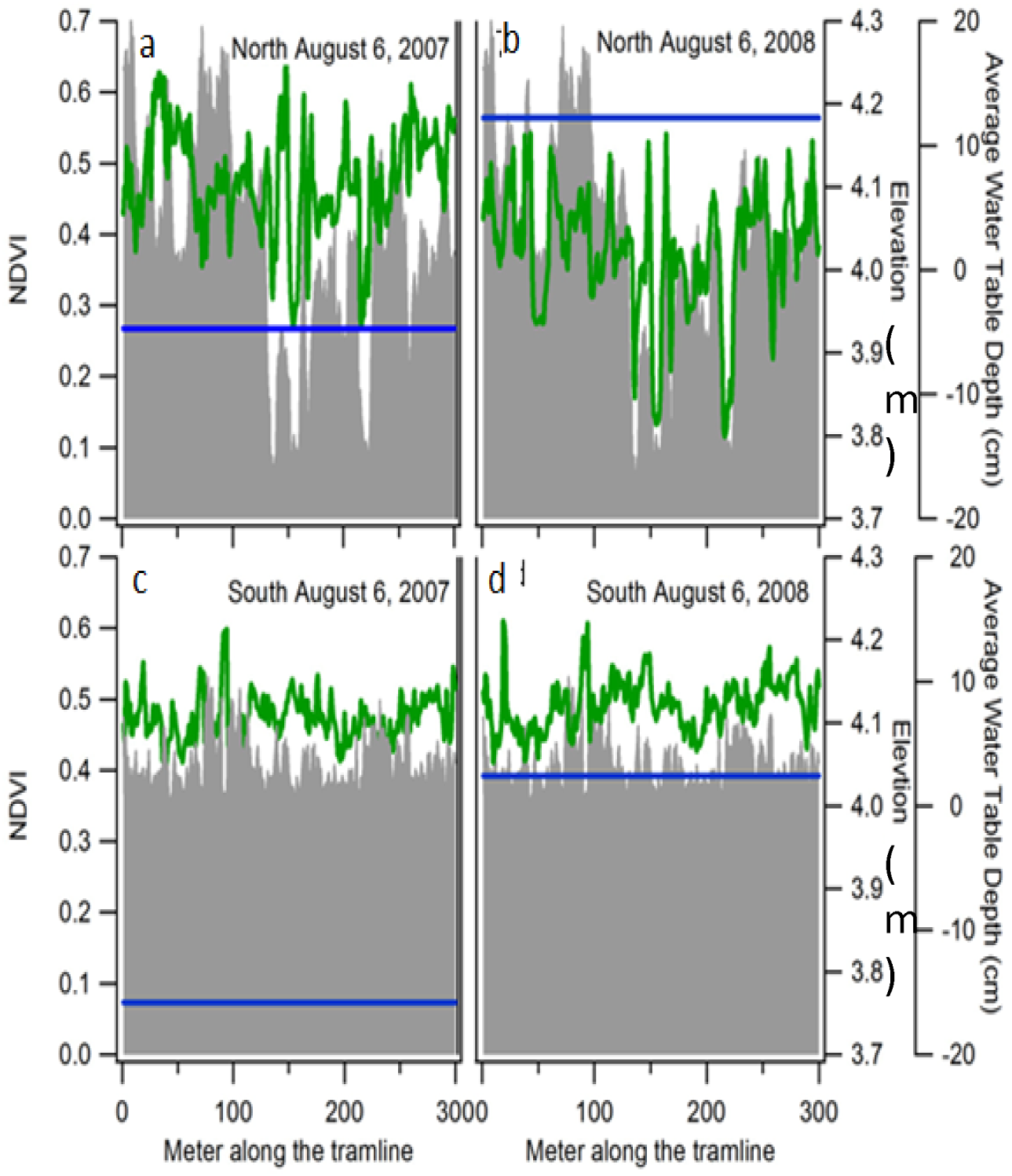
Comparison of modeled (no treatment) vs. measured NDVI and WTD (treatment) time series for the north and control treatments in 2008 and 2009. NDVI and WTD were calculated using the models developed given in figure 4 above for the snow free growing seasons of 2008 and 2009.

**Figure 5.**
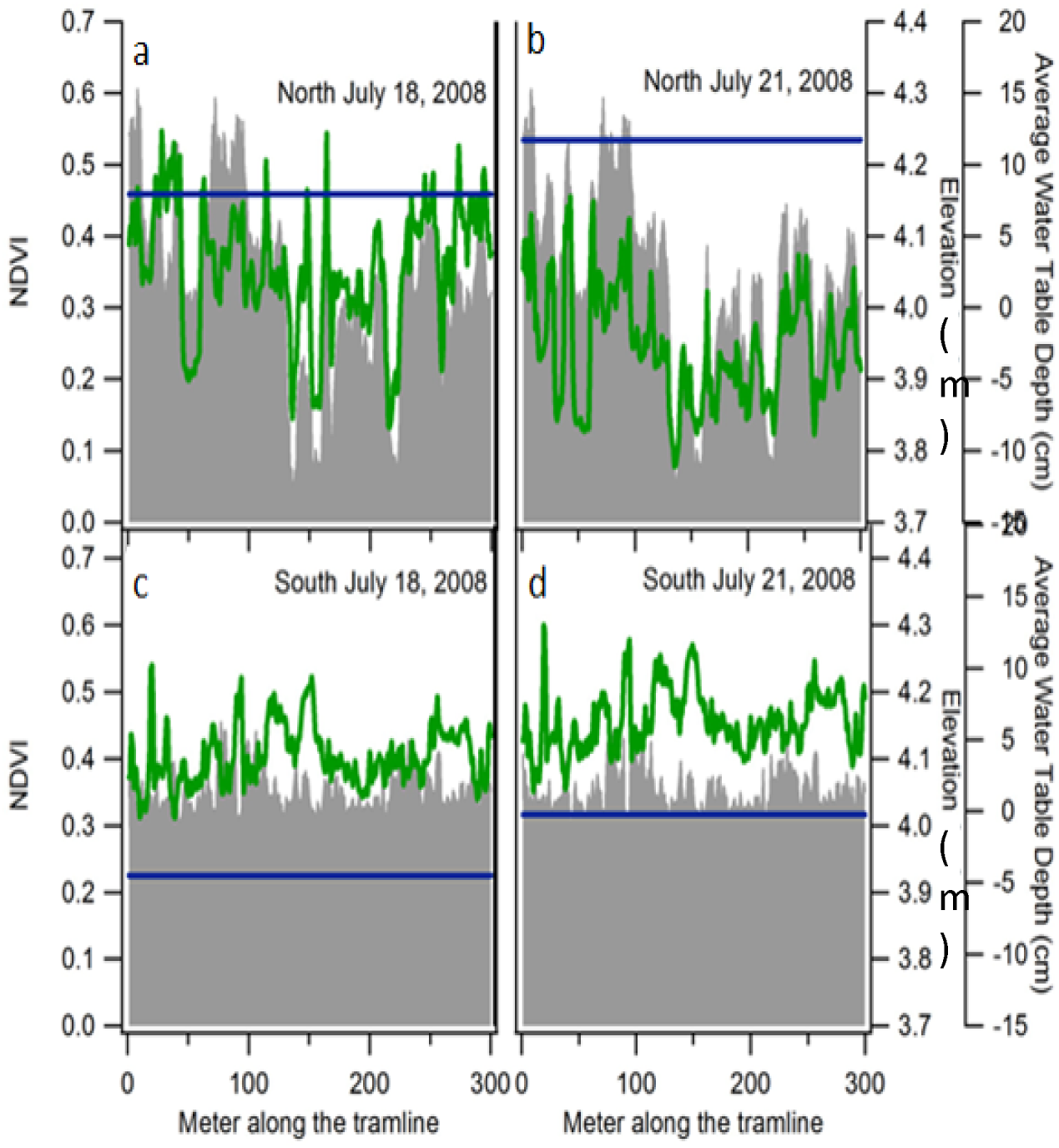
Here the green line indicates NDVI, the blue line indicates average water table depth and the grey line indicates elevation of the tramline area. The above figure shows how a rise in above ground water table depth on the north tramline area lowers the NDVI values on July 21, 2008 shown in 8b compared to the NDVI values for July 18, 2008 shown in 8a. This effect is similar to the effect observed in seasonal patterns of NDVI for north tramline on DOY 210 in figure 5c. Even though below ground water table depth was raised on the south tramline on July 21, 2008 shown in 8d compared to July 18, 2008 shown in 8c, the NDVI values did not decrease.

**Figure 6.**
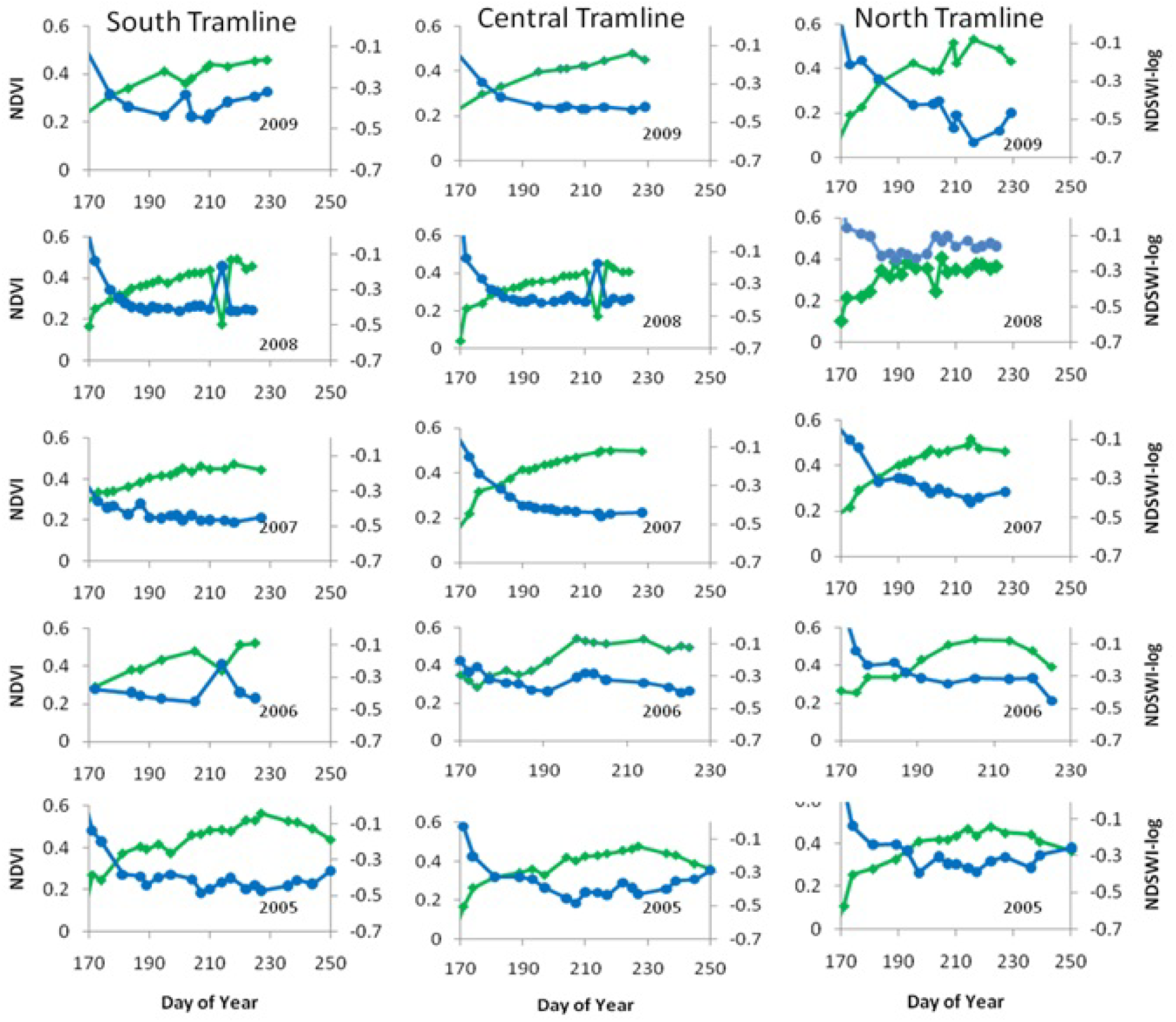
Seasonal dynamics of mean NDVI (green) and NDSWI-log (blue) for all three tramlines 2005 – 2009. 2005 – 2007 were pre-treatment years and 2008 – 2009 were treatment years.

**Figure 7.**
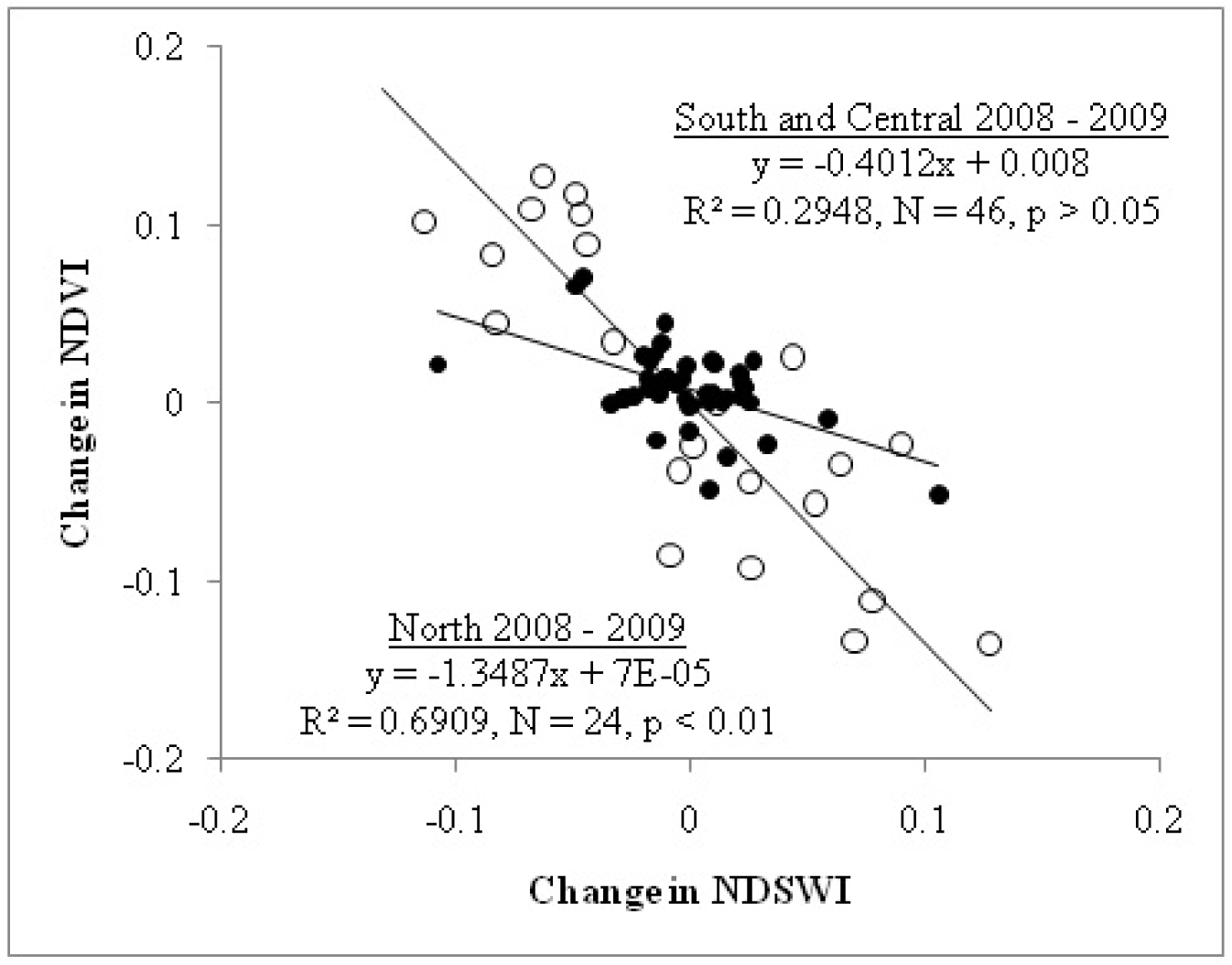
Relationship between the change in NDSWI and NDVI in a measurement period for the north (open circle), and the South and Central tramlines (solid circles) in 2008 and 2009. The north tramline area (flooded) showed a strong correlation indicating that change in surface water affects NDVI values.

**Figure 8.**
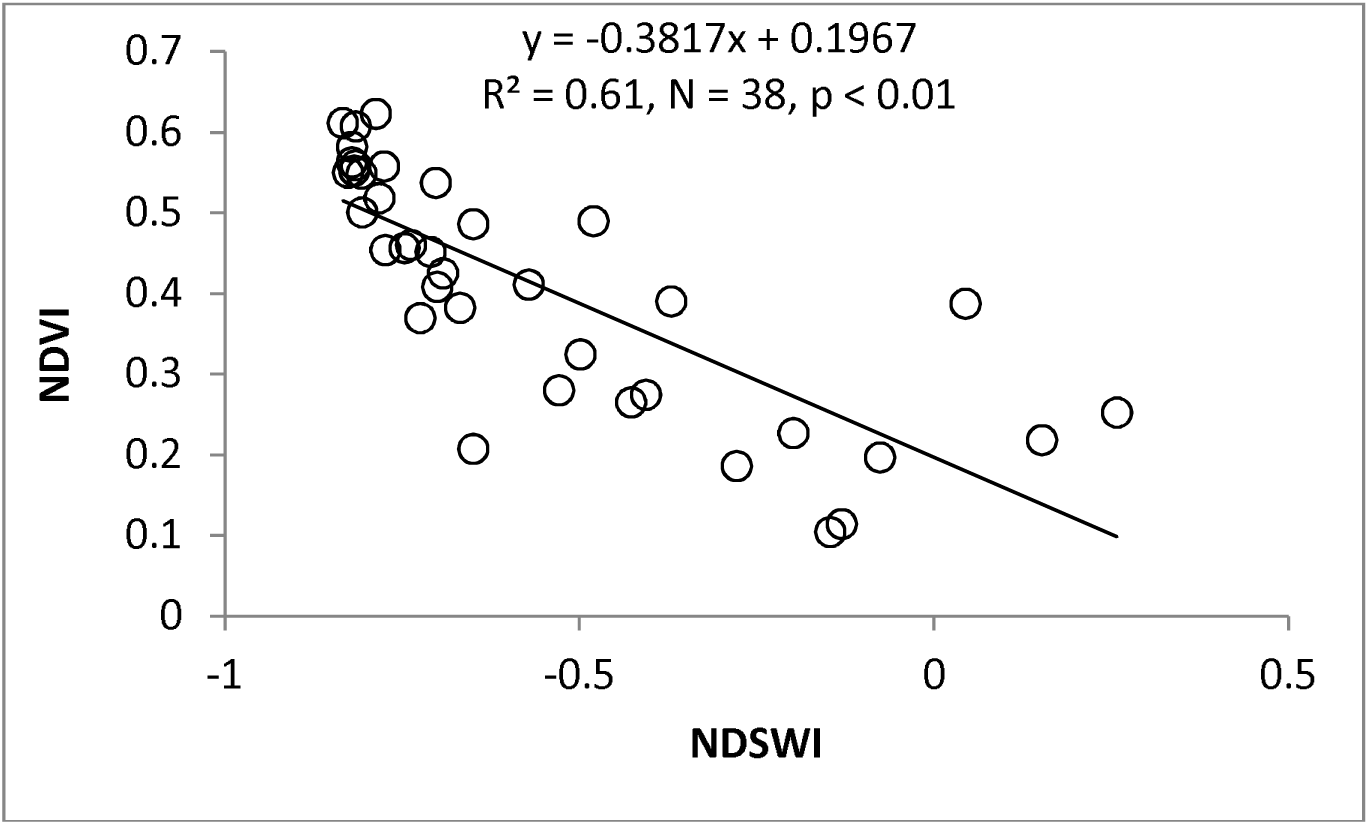
Relationship between NDSWI and NDVI using MODIS 8-day composite time series data for 2000 – 2010 between day 185 and day 297 for each year.

To study how the relationships of NDSWI and NDVI held at the landscape level, 250m pixel size MODIS 8-day composite data were extracted from the Oak Ridge National Laboratory as a 2km by 2km study area using the study site as the center of the subset using data. NDVI values were created using Equation (3.1) and Band 1 (620-670nm ~ Red) and Band 2 (841- 876nm ~ NIR). NDSWI values were calculated using Equation (3.2) and Band 3 (459-479nm ~ BLUE) and Band 5 (1230-1250nm ~ NIR). NDVI and NDSWI values were calculated for the average value of the entire subset swath at a given time period and also for individual pixels within the subset (n = 64 pixels). Regression models were developed between NDSWI and NDVI for the average values of the entire subset and for the individual pixels for the subset to investigate the effect of surface water on NDVI.

## Results and Discussion

### Seasonal and inter-annual variability in land-surface phenology

The transition of the tundra landscape near Barrow as it transitioned from snow-melt in late spring to peak greenness in late July to early August was effectively captured with photographs of site conditions acquired each time the tramlines were sampled. Surface reflectance spectra were also able to capture these transitional states of the landscape throughout the growing season (Fig. 3.1). The snow-melt state of the landscape (Fig. 3.1a), which typically occurs in early June, is represented by the spectrum given in figure 3.1b where the water absorption feature at approximately 970nm is clearly visible. Within the study area and following snow-melt, the landscape is generally flooded (Fig. 3.1c), which is represented by the rather flat reflectance spectra (Fig. 3.1d) and prominent water absorption feature. Following draining, infiltration, evapotranspiration and plant growth (Figs. 3.1e and 3.1g), the spectral signature of vegetation becomes more dominant and no longer includes a strong water absorption feature (Fig. 3.1f). This includes the development of the red edge, which is the rapid change in reflectance in the near infrared range (around 700nm) due to absorption by chlorophyll and a subtle water absorption feature in the 900 nm band through July. Peak growing season typically occurs in early to mid August (Fig. 3.1g) (La Puma *et al.*, 2007). NDVI derived from reflectance spectra given in figure 3.1 (Fig. 3.2), show strong seasonality associated with snow melt and greenup during the typical snow free period in the study area.

Seasonal plots of mean NDVI values for each tramline and year of study showed subtle seasonal differences but no striking differences inter-annually (Figs. 3.3a, 3.3b, and 3.3c). 2006 showed an early snowmelt compared to other years and may in part be related to snow removal associated with the extension of each of the tramlines. The south tramline area had an early snowmelt in 2007. North tramline NDVI values for the 2008 season showed a sudden drop around day 200 that was followed by an increase in NDVI at the next measurement day, after which NDVI decreased again around day 210 and remained steady at a low value for the rest of the measurements made during that snow free period. Similar sudden drops in NDVI values were observed for the central and south tramlines, around day 210 for 2008. This sudden drop in NDVI values for the central and the south tramlines did not follow the patterns observed at the north tramline. NDVI values at the south and central tramlines increased after the sudden drop and followed the same seasonal pattern as described for the north tramline. The short-term variability in NDVI values are investigated in greater detail in figures 3.6 and 3.7 below. Between 2005 and 2009, seasonal peak values of NDVI always occurred earlier for the north tramlines compared to the south and central tramlines. Seasonal peak greening at the North tramline was five days earlier than the central and south tramlines in 2005, ten days earlier in 2006, two days earlier in 2007, and twelve days earlier in 2008, and ten to twelve days earlier in 2009 (Fig. 3.3d).

### Treatment effects on inter-annual land-surface phenology

Correlations between south tramline NDVI values for 2005 – 2007 and NDVI values for this same period at the central and north tramlines showed strong relationships (R^2^ = 0.89, P < 0.001 for south vs. central tramlines figure 3.4a; and R^2^ = 0.88 P < 0.001 for south vs. north tramlines Fig. 3.4b). Similarly, regressions between WTD values for 2007 for south vs. central and south vs. north tramlines showed strong linear relationships (R^2^ = 0.96, P< 0.001 for south vs. central, Fig. 3.4c; and R^2^ = 0.96, P < 0.001 for south vs. north Fig. 3.4d). These strong relationships allowed for the experimental effects on WTD and NDVI to be quantified by calculating the difference between modeled values and measured values for WTD and NDVI during 2008 and 2009.

At the north tramline, measured vs. modeled WTD showed that measured WTD was consistently higher than modeled WTD for the majority of the 2009 growing season (Fig. 3.5a), while the measured WTD was only consistently higher than the modeled values towards the later part of the season starting around day 205 in 2008 (Fig. 3.5c). For both 2008 and 2009, measured WTD was consistently lower than the modeled values for the central tramline (Fig. 3.5b, 3.5d). These suggest that the flooding treatment was most successful during 2009 and that a successful draining treatment was established in both 2008 and 2009.

At the north tramline, measured vs. modeled NDVI showed that measured NDVI was consistently but only slightly higher than modeled NDVI throughout the growing season in 2009 (Fig. 3.5a) while the measured NDVI was slightly lower than the modeled values towards the later part of the season starting around day 205 for 2008 (Fig. 3.5c). For the central tramline, measured NDVI values varied considerably throughout the growing season in both 2008 and 2009 (Fig. 3.5c). Little difference between modeled and measured NDVI was observed in 2008 (Fig. 3.5d) but NDVI was higher than modeled values in 2009 after DOY 195 (Fig 3.5b).

### Relationship between land-surface phenology, microtopography and water table depth

To investigate phenomenon associated with the sudden drop in NDVI values observed in 2008, spatially detailed plots of NDVI, microtopography, and WTD were constructed for the north and south tramlines for measurements made on July 21 and August 6 in both 2007 and 2008. These periods varied considerably in WTD and the response of NDVI between experimental treatments, and thus were used to further investigate the relationship between NDVI and surface hydrology. Intercomparison of these plots (Fig. 3.6) showed that average WTD for the north tramline was approximately 15 cm higher in 2008 than in 2007 and approximately 12 cm above ground level. This meant that areas of lower topography along the north tramline were flooded in 2008 whereas the water table was below ground level on the same day in 2007. Corresponding with this, NDVI values for areas of lower elevation along the north tramline were lower in 2008 than in 2007. The same intercomparison for the south tramline (control) showed that NDVI values were not substantially different between the two years, despite similar absolute differences in mean WTD, which remained below ground level. Mean WTD increased by approximately 5 cm between July 18^th^ and July 21^st^ in 2008 at the north tramline due to flooding treatment. This resulted in a similar response in NDVI to the inter-annual differences described above between 2007 and 2008 and NDVI decreased markedly in the low lying areas (Fig. 3.7). Mean WTD for the south tramline also increased by approximately 5 cm (from -5cm to 0cm, i.e. below ground level) over this same time period but this did not result in a change in NDVI. These investigations suggest that the presence of water above ground level interferes with NDVI signatures over short to long term periods, which is investigated in greater detail in Section 3.6 below.

### Environmental controls of land-surface phenology

The regression tree analysis performed on data from all years (2005-2009) was able to account for approximately 74% of the variability in the dataset (Fig. 3.8). The primary control of NDVI for DSSM ≥ 28 was WTD < 6.39cm. Between DSSM of 18 and 28, WTD ≥ 6.39cm was the primary control of NDVI.

### Relationship between NDSWI-NDVI (surface hydrology and greenness)

Simultaneously plotting NDSWI and NDVI over the course of the snow free growing season showed a strong negative relationship (Fig. 3.9). Low values of NDVI were recorded when high values of NDSWI were also recorded and vice-versa (Fig. 3.9). This effect was particularly prominent at the north tramline during treatment years (2008 and 2009). Linear regression analysis between change in NDSWI and change in NDVI within a sampling period in 2008 and 2009 showed a strong correlation (R^2^=0.69) for the flooded section while this relationship was not so strong for the control and the drained sections (R^2^=0.29) (Fig. 3.10). At this time, WTD was largely below ground in the central and south treatment areas, further suggesting that the measurement NDVI is only affected by dynamic fluctuations in WTD when WTD is below ground level. Linear regression analysis between NDSWI and NDVI calculated from the MODIS 16 day composite time series data for 2000 – 2010 between DOY 185 – 297 showed a R^2^ value of 0.61 indicating that in the relationship between NDSWI and NDVI scales to the satellite scale (Fig. 3.11).

## Discussion

The study of LSP using remote sensing (**Friedl et al. 2006, Henebry et al. 2005**), has seen significant progress over the past two decades (Zhang *et al.*, 2006), (White *et al.*, 2009), (Zhang *et al.*, 2003), (Myneni *et al.*, 1997). Numerous studies have used time series vegetation indices derived from reflectance data acquired by satellite sensors to document and improve understanding of fundamental landscape to continental scale ecosystem properties (Bhatt *et al.* 2010, (Jia *et al.*, 2009), **Running et al. 2008**, (Myneni *et al.*, 1997). LSP is a key indicator of ecosystem dynamics that are susceptible to environmental change (Myneni *et al.*, 1997). Detecting biotic responses such as phenology to a changing environment is essential for understanding the consequences of global change including impacts to ecosystem carbon balance (Merbold *et al.*, 2009), (Wolf *et al.*, 2008) energy balance (Euskirchen *et al.*, 2007), (Chapin *et al.*, 2005), and biodiversity **(Foster et al. 2010)**. This study aimed to monitor seasonal and inter-annual trends in LSP and understand how LSP responds to altered surface hydrology and inter-annual climatic and other environmental variability using reflectance data from the robotic tram system. The robotic tram system used in the study provided an excellent platform to make continuous and repeatable ground-based measurement of LSP over the study period of 2005-2009 as part of the biocomplexity experiment.

We expected to observe substantial changes in LSP in response to the flooding and draining experiment. Instead, we observed a response to the experimental treatments that had a similar magnitude of variability to that associated with inter-annual variability. In retrospect, the LSP results are expected now that more detail on the inter-annual variability in WTD has been established. This study has shown that inter-annual variability in WTD is just as great as experimental effects associated with the treatments, hence the treatments invoke realistic environmental states and serve to extend the opportunity to observe condition states that are plausible for the study site. Experimental flooding treatments in 2008 were not as consistent as those in 2009. The year 2008 was wetter and the capacity to flood was limited by the experimental infrastructure and natural topography which did not permit flooding beyond the level studied as water simply poured out over the top of the topographic margins intended to contain the treatment. Similar problems were minimal in 2009. These limitations in flooding treatments were visible in the LSP patterns for 2008 and 2009 as 2008 showed more fluctuations in the seasonal pattern of LSP than in 2009.

The 2006 seasonal pattern of NDVI showed the earliest snow-melt and highest peak season values for the control and the drying treatment areas. This was probably due to snow being removed in early spring for construction that extended the tramlines by 100m to the east of each of the tramlines for an extension of the tramlines. This finding supports previous finding that early snowmelt can result in higher peak season NDVI and productivity (La Puma *et al.*, 2007), (Kimball *et al.*, 2006).

The strong correlation between the south and central and north tramlines for NDVI and WTD indicated that values for the central and north tramlines can be modeled from the south tramline to assess the likely experimental impact of the draining and flooding experiments respectively. This was important for this experiment because it was unreplicated, and the treatments were held relative to inter-annual variability such that flooding during a dry year presents WTD’s similar to what is experienced in the control treatment in a wet year. There was no marked difference in landscape phenology associated with the treatments. This finding is similar to those reported by (Olivas *et al.*, 2011) and (Zona *et al.*, 2011) for CO2 flux. This is unlike findings reported for thaw depth where deeper thaw depths were reported for 2008 and 2009 for the flooding treatment area compared to 2006 and 2007 (Shiklomanov *et al.*, 2010).

The effect of the treatment on LSP was more subtle than expected. To better understand factors controlling the spatiotemporal variability in the LSP patterns of the three treatment areas, regression tree analysis was used to analyze the interplay between NDVI and a range of factors shown by other studies to influence LSP and other ecosystem properties and processes. These included DSSM, daily average temperature, daily total PAR, and WTD for snow-free period of the growing season. Regression tree analyses do not allow for derivation of causation. Instead they are intended for the data mining and exploratory analyses that facilitate the discovery of tipping points and correlations with multiple and co-occurring environmental drivers. Here regression trees suggest that the spatiotemporal variability in NDVI was strongly controlled by DSSM and WTD thereby giving us new understanding of the environmental controls of NDVI spatiotemporal variability. We have known DSSM to be an important factor controlling seasonal NDVI but this appears to be the first ground based study in the arctic to show that variable surface hydrology can impact capacities to adequately determine NDVI.

More detailed analyses of LSP along tramlines on specific dates of measurement indicate both the challenge of acquiring NDVI in the presence of surface water strongly interferes with spectra used to derive NDVI. In this detailed study, the importance of micro-topography in influencing NDVI and surface water coverage was strongly apparent. Other studies have also shown how microtopographic variability can control ecosystem properties and processes such as species distribution **(Henry 1998)**, soil moisture and WTD, surface energy budgets, trace gas flux (**Gamon et al. in press**, (Zona *et al.*, 2011). Analysis for pre/post treatment variability and for short-term inter-annual variability showed that fluctuations in WTD and surface water cover had the greatest impact in low lying areas and NDVI decreased as WTD and surface water coverage increased. This phenomenon was observed between years, within a season and with environmental variability that was experienced over just a few days. Although there is likely to be some variability in the above ground green plant biomass that is measured as NDVI in such measurements, the majority of the variability experienced can be explained by surface water coverage, which appeared to be well described by NDSWI, a spectral index for estimating surface water cover and depth (Goswami *et al.*, 2011).

The statistically strong relationship between NDSWI and NDVI in the MODIS 2km by 2km subset created for the study area also indicated that surface water could play an important role in controlling NDVI values at the satellite scales. The implications of this finding is important – especially for large scale studies in the Arctic that use satellite data where NDVI is used as an indicator of greening (Bhatt *et al.*, 2010), (Jia *et al.*, 2009). Results from this study suggest that WTD is near the ground surface, greening can be documented as a result of drying.

## Conclusions

This study characterized seasonal and inter-annual trends in land-surface phenology in an arctic tundra landscape for a period of five years and included measurement of pre and post flooding and draining treatment conditions in a large scale experimental manipulation. To our knowledge, no studies have reported such an extensive spatio-temporal dynamics of land-surface phenology in an arctic landscape using near surface remote sensing techniques. The overarching goal of the study was to further understand the spatio-temporal dynamics and controls of LSP in an arctic tundra landscape using optical remote sensing and investigate if altered surface hydrology could impact LSP. We also sought to investigate these effects on data collected from ground and satellite platforms. Findings indicate there were no major differences between inter-annual patterns or treatment differences in land-surface phenology in the study area. There were some abrupt changes in intraseasonal patterns, which seemed to occur due to changes in surface hydrology as a result of experimental treatments and weather events such as snowfall events. A relatively high spatial resolution analyses also suggested that changes in land-surface phenology appeared to be related to changes in surface hydrology in areas of low lying microtopography where WTD was above ground. NDSWI and NDVI showed a strong negative relationship when water table was above ground level indicating that an increase in the cover of surface water, which is measureable with NDSWI, decreases NDVI and vice versa. These relationships held both at the high resolution tramline scale (plot level) and at the landscape level measured from reflectance derived from a globally orbiting satellite platform. Although widespread greening of the Arctic has been documented, this study suggests that if there has been a drying of landscapes similar to surface that of the study area, NDVI could have increased, not necessarily as a result of increased green plant biomass but because there is less water, which distorts accurate measurement of NDVI. Such drying of arctic landscapes has important implications for further understanding arctic ecosystem change and predicting the future state of the Arctic and Earth Systems.

## Acknowledgements

This project was supported by the U.S. National Science Foundation (ASSP-0421588). We are grateful to the Ukpeaġvik Iñupiat Corporation (UIC) for permitting the Biocomplexity experiment on the Barrow Environmental Observatory and the Barrow Arctic Science Consortium and CH2M Hill Polar Services (formerly VECO Polar Services) for logistical support, construction services, and ongoing tramline maintenance.

